# A robust, reproducible, accessible and scalable protocol for generating three-dimensional human gastruloids

**DOI:** 10.64898/2026.05.23.727089

**Authors:** Arghakusum Das, Shekhar Patil, Karthik Ravi, Maneesha S Inamdar

**Affiliations:** Jawaharlal Nehru Centre for Advanced Scientific Research, Bengaluru, Karnataka 560064, India; Institute for Stem Cell Science and Regenerative Medicine GKVK – Post, Bellary Road, Bangalore 560065, Karnataka, India

**Keywords:** human gastruloids, human embryonic stem cells, anteroposterior axis, gastrulation, robust & reproducible protocol, single-cell RNA sequencing, BJNhem20, BJNhem19, Indian hPSC lines

## Abstract

The rapid rise of stem cell-based human embryo models has reignited interest in studying early human development while offering a promising platform to de-risk drugs. Among these, three-dimensional human gastruloids provide a tractable system to model symmetry breaking, germ layer specification and axial organization. However, existing gastruloid protocols remain expensive, specialized, variable and evaluated in a limited number of human pluripotent stem cell (hPSC) lines, restricting broader adoption. Here, we present a simple, robust, standardized gastruloid protocol achieving greater than 90% elongation efficiency with low inter- and intra-experimental variability, developed primarily in BJNhem20, a well-characterized Indian-origin human embryonic stem cell line. Further, we show that the protocol is applicable in a diverse set of hPSC lines. Using a TBXT (Brachyury)-GFP reporter in BJNhem20, we optimized cell seeding density, induction medium and Wnt activation strength, guided by real-time, quantitative assessment of mesoderm induction and symmetry breaking, allowing precise titration of CHIR99021. Comparative testing identified an in-house “Essential 6” medium formulation as the most consistent condition for robust TBXT induction. Optimization of aggregation density produced reproducible gastruloids with polarized TBXT expression and consistent axial elongation, within 72 hours. Single-cell RNA sequencing of individual gastruloids confirmed high transcriptional reproducibility and conserved lineage clusters, aligned with developmental trajectories. Cell line-specific CHIR99021 titration was sufficient to successfully transfer the optimized protocol to two additional lines, BJNhem19 and RUES2-GLR. This simplified and robust protocol reduces costs and improves accessibility, enabling broader application of stem cell-based human embryo models.

## Introduction

Human gastrulation occurs post-implantation, in week 3 of embryogenesis, during which the body axes are established, and the three primary germ layers are specified and spatially organized. Gastrulation also falls within the embryonic period of greatest sensitivity to teratogenic exposure and is the developmental stage at which many structural congenital malformations originate (Herion et al., 2014; Mantziou et al., 2021; Obićan and Krstić, 2025). Despite its central importance, direct investigation of human gastrulation remains extraordinarily challenging due to ethical constraints and the limited availability of post-implantation material.

Three-dimensional (3D) human gastruloids developed by Moris et al. in 2020, are aggregates of defined numbers of pluripotent stem cells that under controlled culture conditions, undergo symmetry breaking, anteroposterior axis formation and spatiotemporally organized specification of derivatives from all three germ layers. Human gastruloids correspond to approximately Carnegie stage 8–9 of human development, encompassing features of 19–21-day-old embryos, including early signatures of somitogenesis (Moris et al., 2020). The self-organizing nature of gastruloids, their scalability and compatibility with live imaging, chemical perturbation and single-cell analysis make them a uniquely powerful platform for studying human developmental processes.

However, the adoption of human gastruloids beyond a small number of specialist laboratories remains limited by practical barriers. Successful generation requires careful optimization of three sequential steps: the initial cell state, a timed CHIR99021 pre-treatment to activate canonical Wnt/β-catenin signaling and the subsequent aggregation and culture of defined cell numbers (Moris et al., 2020; Moris et al., Protocol Exchange, 2020). Each step is highly sensitive to initial culture conditions, including founding cell number, culture medium composition, morphogen dosage and reagent quality (Moris et al., Protocol Exchange, 2020; Hamazaki et al., 2024; Heemskerk et al., 2019).

Robustness and reproducibility of gastruloids, both within and between experiments and cell lines, remain essential for broader adoption in developmental modeling and applications such as drug safety assessment and systematic perturbation studies. However, inter-experimental variability, a long-standing challenge across hPSC-based systems (Volpato & Webber 2020), is particularly pronounced in gastruloid generation, where small fluctuations in initial conditions are amplified by the self-organising dynamics of the model. As with hPSC differentiation more generally, human gastruloids require empirical optimization for individual hPSC lines. The protocol has been successfully established for H9, RUES2-GLR and MasterShef7 hESC lines and WTC-11 and NCRM1 hiPSCs and their derived reporter cell lines (Moris et al., 2020; Hamazaki et al., 2024; Makwana et al., 2025).Nevertheless, the combined impact of the limitations—protocol sensitivity, restricted crossline validation and the absence of a live readout for real-time monitoring of induction quality- has constrained the accessibility and scalability of the human 3D gastruloid model.

A further, underappreciated dimension of this problem lies in the limited genetic diversity of hPSC lines currently used in gastruloid research. Genetic variation between populations affects disease susceptibility, drug-metabolising enzyme function and the predictive power of clinical genetic risk scores (Sirugo et al., 2019; Martin et al., 2019; Zhou et al., 2017). These would remain largely untested if a limited number of cell lines were used for drug testing or disease-based screening. As gastruloids advance toward applications in personalized medicine, teratogenicity screening and cross-population developmental studies, this gap limits the generalizability of findings derived from the current set of reference lines.

Here, we present a simple, scalable and standardized framework for generating human 3D gastruloids with near 100% efficiency and substantially reduced inter-experimental variability. Using a TBXT-2A-EGFP knock-in reporter in the well-characterized Indian-origin hESC line BJNhem20 (Inamdar et al., 2009; Patil et al., 2026) as the primary model system, we identified key parameters governing gastruloid induction and successful formation. By real-time, quantitative monitoring of mesoderm specification and symmetry breaking, we show that initial culture medium, CHIR99021 concentration and *TBXT* expression status are critical parameters to control for successful gastruloids generation. Our optimized protocol reduces costs and increases efficiency of gastruloid formation and is applicable to multiple genetically distinct hPSC lines. This robust and accessible platform lowers technical barriers, enables broader adoption across diverse research settings and expands the utility of the gastruloid model for translational applications.

## Materials and Methods

### hPSC cell lines, culture and ethics statement

hPSC lines used in this study included the well-characterized Indian-origin hESC lines BJNhem20 and BJNhem19 (Inamdar et al., 2009; ISCI, 2011) that have been used in multiple studies and the germ layer reporter RUES2-GLR that has been widely used for gastruloid generation (Martyn et al., 2018; Moris et al., 2020). BJNhem19 and BJNhem20 are available from (https://scvbljncasr.wixsite.com/scvbl/resources#anchors-lv7w62xc3) and have also been deposited in the UK Stem Cell Bank (UKSCB accession nos. R-08-021 and R-08-022), are listed on hPSCReg (JNCSRe001-A and JNCSRe002-A) and the NIH Registry (NIH Registration no. 0083 and 0084). A TBXT-2A-EGFP knock-in reporter was introduced into both BJNhem20 (Patil, Das and Inamdar, 2026) and BJNhem19 (JNCSRe001-A-1) by CRISPR-Cas9 gene targeting and validated prior to use; these lines are referred to throughout as BJNhem20 TBXT-EGFP and BJNhem19 TBXT-EGFP respectively. All lines were maintained under feeder-free conditions in mTeSR Plus medium (STEMCELL Technologies) on Matrigel (Corning) coated plates at 37 °C with 5% CO_2_ and medium was replaced every 48 hours. Colonies were passaged every 3–5 days using mechanical dissociation. Cells were routinely monitored for morphology and confirmed to be mycoplasma-negative at regular intervals. All hPSC work was conducted after obtaining approval from the Institutional Committee for Stem Cell Research (approval no. 8/ICSCR/XVI/MI) in accordance with the National Guidelines for Stem Cell Research, Govt. of India.

### Preparation of in-house Essential 6 medium (iE6)

Essential 6 (E6) medium was prepared in-house from individual components deduced based on the Essential 8 formulation described by Chen et al. (2011), with omission of transforming growth factor beta 1 and fibroblast growth factor 2. The following components were combined in Dulbecco’s Modified Eagle Medium/F12 (1:1, Invitrogen): sodium bicarbonate (Sigma-Aldrich), L-ascorbic acid 2-phosphate (Sigma), sodium selenite (Sigma), human transferrin (Roche) and insulin (Roche). All components were used at concentrations specified by Chen et al. (2011). The medium was filter-sterilized through a 0.22 µm membrane, aliquoted into airtight containers and stored at 4 °C. Each batch was prepared fresh and used within 1–2 weeks of preparation. Medium was handled on ice.

### Cell plating density and CHIR99021 pre-treatment

Cells were dissociated into single-cell suspensions using TrypLE Express (Thermo Fisher Scientific) and seeded onto Matrigel-coated 35 mm tissue culture dishes in mTeSR Plus medium supplemented with 10 µM Y-27632 (ROCK inhibitor; Sigma). Multiple seeding densities were tested, between 20k and 100k to identify the optimal density. Cells were allowed to attach and recover for 24 hours, after which the medium was replaced with iE6 medium containing required dose of CHIR99021 (Tocris Biosciences; dissolved in DMSO as a 10 mM stock). Pre-treatment was carried out for 24 hours at 37 °C with 5% CO_2_. The optimal pre-treatment concentration for each line was established empirically based on TBXT-GFP expression. Each new reagent lot was calibrated prior to use in gastruloid experiments.

### Cell aggregation

Following CHIR99021 pre-treatment, cells were washed once with phosphate-buffered saline (PBS) and dissociated into single-cell suspensions using TrypLE Express. Cells were counted using a hemocytometer and resuspended in iE6 medium supplemented with 10 µM Y-27632 and CHIR99021. Multiple cell numbers between 200-1000 cells were tested by adding the cells in 40 µL of iE6 medium into non-adherent, U-bottom 96-well plates (Greiner Bio-One) followed by centrifugation at 70 × g for two minutes and 170 × g for an additional two minutes, to facilitate aggregate compaction and identify the optimal number for gastruloid formation.

### Induction of gastruloid patterning

At the time of aggregation (day 0), CHIR99021 pulse concentrations between 0.5 µM and 3.5 µM were tested for each line. CHIR99021 was diluted at the first medium change 24 hours after induction (day 1), by adding 150 µL of fresh iE6 medium. Subsequently the medium was replaced daily with 150 µL of fresh iE6 throughout the culture period. Gastruloids were cultured for 72 hours post-aggregation unless otherwise stated. All aggregates were maintained at 37 °C with 5% CO_2_.

### Imaging and Morphometric analysis of gastruloids

Phase-contrast and fluorescence images of gastruloids were acquired every 24 hours post-aggregation using an EVOS M7000 imaging system (Thermo Fisher Scientific) at 10× magnification. The entire 96 well plate was scanned simultaneously in bright field and in 488 channel to detect GFP. Images were exported and analyzed using a custom MATLAB script (MATLAB R2023a, MathWorks) adapted from Moris et al., 2020. Gastruloid outlines were segmented using a polygon-based manual annotation tool and the following morphological parameters were extracted for each structure: projected area, perimeter, circularity (defined as 4π × area / perimeter^2^) and aspect ratio (major axis length / minor axis length) (Supplementary Figure S2). Circularity range is from 0 (infinitely elongated) to 1 (perfect circle), whereas aspect ratio increases with elongation.

### Immunofluorescence staining

Gastruloids were collected at 72 hours post-aggregation, transferred to 24-well plates and washed once in PBS. Samples were fixed in 4% paraformaldehyde in PBS for 30 minutes to one hour at room temperature. Fixed gastruloids were washed three times in PBS and then permeabilized and blocked overnight at 4 °C in 1× PBSFT (PBS containing 0.2% Triton X-100 and 10% fetal bovine serum). Primary antibodies were applied in 1× PBSFT and incubated overnight at 4 °C on an orbital shaker. Following primary antibody incubation, samples were washed three times in 1× PBSFT (10 minutes per wash) and incubated with fluorescent secondary antibodies for one hour at room temperature or overnight at 4 °C in the dark. Nuclei were counterstained with Hoechst 33342 for 5–10 minutes at room temperature. Gastruloids were washed three times in PBS and mounted in 70% glycerol on glass slides. Images were processed and assembled using ImageJ/FIJI (Schindelin et al., 2012).

### Single-cell RNA sequencing

#### Library preparation

BJNhem20 TBXT-EGFP gastruloids were harvested at 72 hours post-aggregation. Four scRNA-seq libraries were prepared: three from individual gastruloids (Dish1_E5, Dish2_B5and Dish3_B10), each selected from a separate, independently established 96-well plate seeded from distinct 35 mm pre-treatment dishes and one from a pooled sample containing three gastruloids drawn from all three independent experiments. Individual gastruloids were selected based on characteristic elongated morphology at 72 hours.

Gastruloids were dissociated into single-cell suspensions by incubation in TrypLE Express for 2–3 minutes at 37 °C with gentle agitation. Dissociated material was filtered through a 40 µm cell strainer and washed in 1× Dulbecco’s PBS. Cell viability was assessed by propidium iodide exclusion staining and only samples with viability exceeding 80% were used for library preparation. Single-cell libraries were prepared using the 10× Genomics Chromium Single Cell 3′ Reagent Kit v3.1 according to the manufacturer’s instructions. Libraries were sequenced on an Illumina NovaSeq 6000 to a target depth of approximately 200 million reads per sample.

#### Computational analysis

Raw sequencing reads were demultiplexed and aligned to the human reference genome (GRCh38) using Cell Ranger (10× Genomics). Gene expression data were normalized using SCTransform (Hafemeister and Satija, 2019), which models technical variation related to sequencing depth and stabilizes variance across genes, following the analytical framework described by Hamazaki et al. (2024). The three individual gastruloid libraries (Dish1_E5, Dish2_B5, Dish3_B10) were integrated using Harmony (Korsunsky et al., 2019) to correct sample-of-origin batch effects. The pooled library was processed independently and was not included in the Harmony integration. Dimensionality reduction was performed using principal component analysis, followed by UMAP (McInnes et al., 2018). Unsupervised clustering was performed using Seurat FindNeighbors and FindClusters (Stuart et al., 2019), yielding nine clusters (clusters 0–8).

Cell type annotation was assigned based on canonical marker gene expression and reference-based classification. Reference label transfer was performed using SingleR (Aran et al., 2019) with the human gastruloid dataset from Hamazaki et al. (2024) as the reference. Manual review of marker gene expression was used to confirm and refine annotations. The final annotated populations were PSM, Cycling_PSM, Somite, Cycling_Somite, Lateral_Plate_Mesoderm, Neural_progenitor, NMP, Stress_response and Unassigned.

### Statistical analysis

Morphometric parameters (circularity and aspect ratio) were compared between CHIR99021 treatment conditions and the DMSO vehicle control using Tukey’s post hoc multiple comparisons test. Statistical analyses were performed in GraphPad Prism. A p-value threshold of p < 0.0001 was used to define statistical significance, as indicated in the relevant figures. No intergroup comparisons between CHIR99021 concentrations were performed; all comparisons were made against the DMSO vehicle control only. Each experiment was one 96 well plate. Measurements were obtained from gastruloids across at least three independent experiments. Comparisons of morphometric parameters were performed between CHIR99021-treated conditions and the DMSO vehicle control; intergroup comparisons between CHIR99021 concentrations were not conducted.

## Results

### Generation of gastruloids in BJNhem20 hPSC line

To address the absence of validated gastruloid protocols in genetically diverse hPSC lines, we aimed to generate gastruloids in BJNhem20, an Indian-origin hESC line (Inamdar et al., 2009). In the protocol developed by Moris et al. (2020) hPSCs are pre-treated with a 24 h CHIR99021 pulse in Nutristem XF, dissociated to single cells, seeded into U-bottomed 96-well plates at a defined cell number in Essential 6 (E6) medium, and then exposed to a second lower concentration CHIR99021 pulse from 24 to 48 h during aggregation. Gastruloids that have broken symmetry and elongated are scored from 72 h onwards (Figure 1A). Adapting this protocol to BJNhem20 required empirical standardization of four parameters: the pre-treatment medium and CHIR99021 dose, the CHIR99021 pulse dose at aggregation, the cell number per well and the aggregation medium itself. Throughout this work, we distinguish and report two levels of reproducibility — intra-experimental, defined as inter-well consistency in aggregate progression within a single 96-well plate, and inter-experimental, defined as consistency in outcomes between independent plates run on different days.

**Figure 1.**
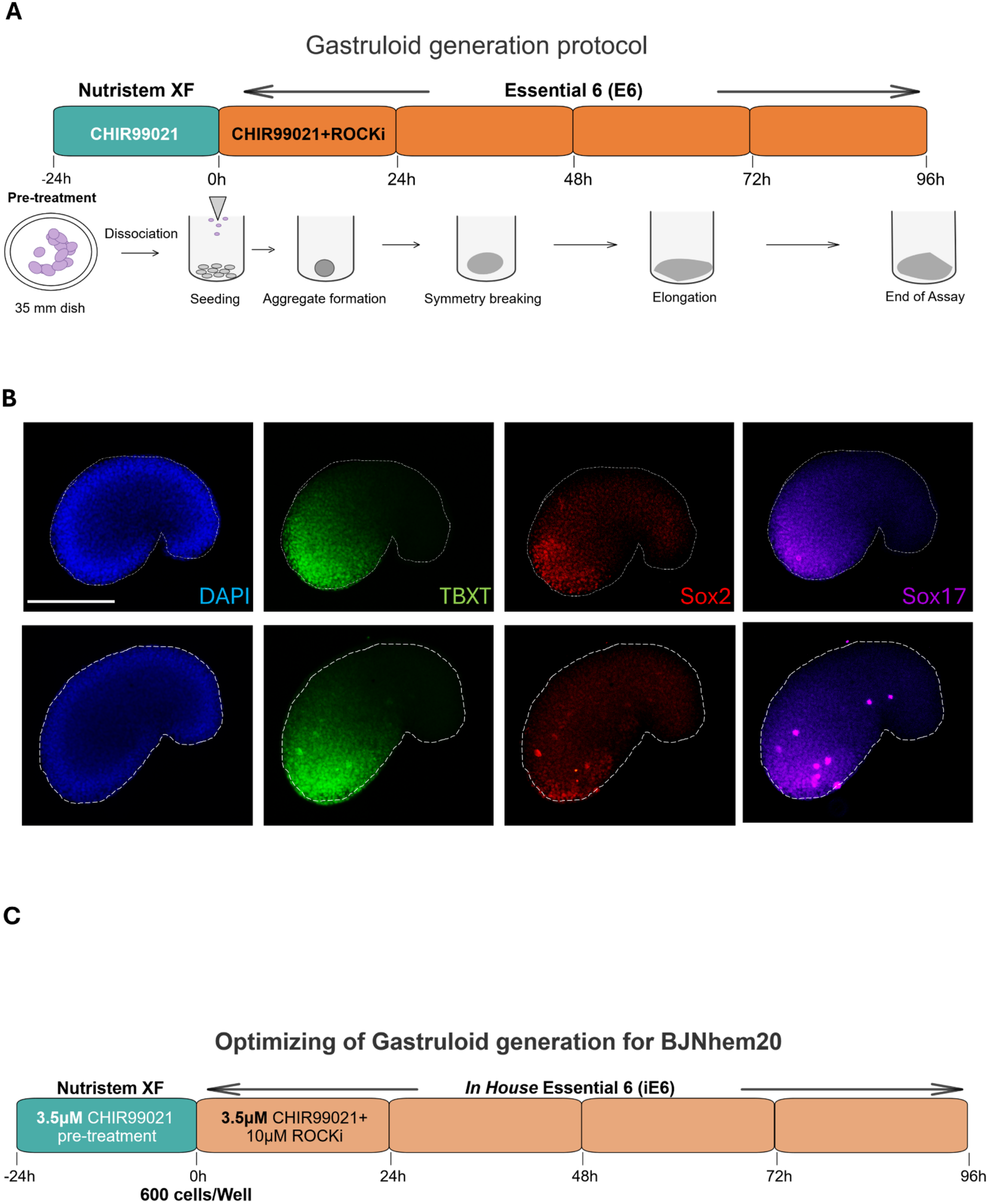
Establishing a gastruloid protocol in BJNhem20 hESC line. (A) Schematic representation of the human gastruloid generation protocol devised by Moris et al. (2020). hPSCs maintained in 2D culture were pre-treated with CHIR99021 in Nutristem XF medium for 24 h (−24 to 0 h), then dissociated to a single-cell suspension and seeded into U-bottomed 96-well plates in commercial Essential 6 (E6) medium. A second CHIR99021 pulse, supplemented with ROCK inhibitor (Y-27632), was applied from 0 to 24 h, after which cells were maintained in E6 medium. Aggregates were imaged at 24 h intervals; symmetry breaking and axial elongation were typically observed between 48 and 72 h post-aggregation. (B) Whole-mount immunofluorescence images of two representative BJNhem20 gastruloids formed in the 3.5 µM CHIR99021 pre-treatment condition, at Day 3 (72 hours) post-aggregation showing expression of TBXT (Green), Sox2 (Red) and Sox17 (Magenta). Nuclei stained with DAPI (Blue). Scale bar: 275 µm. (C) Schematic of the optimised gastruloid generation protocol for BJNhem20. Pre-treatment was performed with 3.5 µM CHIR99021 in Nutristem XF for 24 h. Cells were dissociated and 600 cells per well were seeded into U-bottomed 96-well plate in in-house Essential 6 medium (iE6), with a 3.5 µM CHIR99021 + 10 µM ROCK inhibitor pulse applied from 0 to 24 h post-aggregation. Aggregates were maintained in iE6 thereafter. Morphometric assessment was performed at 72 h, the timepoint at which gastruloid elongation plateaus.

### Optimization of Chiron pre-treatment and pulse dose

For pre-treatment of BJNhem20 in Nutristem XF, we compared 3.5 µM and 5.0 µM CHIR99021 exposure, scoring outcomes by aggregate morphology every 24 h (size, symmetry breaking and elongation) and by spatially restricted germ layer marker expression (TBXT, SOX2 and SOX17) at 72 h (Figure 1B). Aggregates formed under both conditions, but with distinct outcomes. Pre-treatment with 5.0 µM CHIR99021 yielded compact aggregates that failed to break symmetry in all 192 wells, across 2 independent plates (0/96; 0/96 wells). Pre-treatment with 3.5 µM CHIR99021 produced gastruloids in 3 of 5 plates. Interestingly, in experiments that worked, all aggregates that were set up (36/36; 12/12; 84/84) underwent symmetry breaking and axial elongation and acquired the expected marker expression by 72 h (100% intra-experimental reproducibility). In the remaining 2 plates (0/36;0/12 aggregates), aggregates broke symmetry but showed no detectable expression of TBXT, indicating a failure of lineage specification despite morphological progression. This indicated that while BJNhem20 gastruloids can be formed with a 3.5 µM pre-treatment dose, other variables impact gastruloids success. Hence, we adopted 3.5 µM CHIR99021 in Nutristem XF as the pre-treatment condition for BJNhem20.

A CHIR99021 pulse is applied at aggregation to initiate self-organization. Testing 1.5, 2.5, 3.0 and 3.5 µM doses of CHIR99021, we found that all aggregates were spherical at 24 h. By 48 h, 1.5 µM pulsed aggregates remained indistinguishable from DMSO controls, whereas higher concentrations promoted symmetry breaking. By days 3–4, gastruloids in 2.5 µM showed modest elongation with protrusions and uneven edges, compared to 3.0 and 3.5 µM pulse that yielded well-elongated gastruloids with two morphologically distinct poles (Supplementary Figure S1A-B). Quantitative morphometric analysis confirmed this hierarchy in terms of circularity and aspect ratio (Supplementary Figure S1C). Whole-mount immunofluorescence showed the characteristic spatially restricted co-expression of TBXT, SOX2 and SOX17 at the posterior pole (Figure 1C) and was reproducible across at least two independent experiments (Supplementary Table S3). Aggregate length did not increase appreciably between 72 and 96 h under any condition, in agreement with Moris et al. (2020). As 3.0 and 3.5 µM produced equivalent morphometric outcomes in BJNhem20, 3.5 µM was adopted as the pulse range for subsequent experiments. Analyses were from 72 h post-aggregation.

#### Optimization of cell number

To identify the cell number that produced consistent aggregate morphology and elongation, we tested 400, 600, 800 and 1,000 cells per well, drawing on the range previously reported (Moris et al., 2020). At 800 and 1,000 cells per well, aggregates were well-compacted but consistently failed to undergo axial elongation. At 400 and 600 cells per well, aggregates broke symmetry and elongated; 600 cells per well produced the most consistent compaction and morphology across wells and was therefore adopted for all subsequent experiments (Supplementary Table S1).

#### Establishing a reliable aggregation medium: in-house E6 (iE6)

Despite the above standardized parameters, inter-experimental reproducibility at the aggregation step remained poor. Across 24 independent 96 well plates seeded with BJNhem20 cells in commercial E6 medium under otherwise identical conditions, cells formed compact aggregates in only 5 plates (21%); in the remaining 19 plates, cells either failed to aggregate and remained as dispersed single cells or loose, non-cohesive clusters even 24 h post-seeding, or formed aggregates that disintegrated within 48 h. This outcome was independent of the batch number of E6 and was not predicted by visible reagent quality. We reasoned that reagent quality may be compromised during shipping or transit, as is often the case with imported reagents (Inamdar 2023). To remove this dependency, we prepared E6 medium in-house from individually sourced components (iE6), by deducing the formulation from Chen et al. (2011) (see Methods). With iE6 in place of commercial E6 as the aggregation medium, aggregation proceeded consistently across 20 of 20 independent plates with all wells producing well-formed, compact aggregates that did not disintegrate. iE6 was therefore adopted as the standard aggregation medium for all subsequent experiments.

iE6 reliably resolved the aggregation-step failures observed with commercial E6 but did not eliminate variability at the subsequent gastruloid-formation step. Across the same 20 plates, aggregates formed in all wells but only 9 plates produced elongated gastruloids by 72 h, with intra-plate success ranging from 19 of 48 wells to 96 of 96 wells (40–100% per plate; Supplementary Table S2). The remaining 11 plates produced no elongated gastruloids: aggregates formed normally but remained spherical through day 3, indicating that the failure mode had shifted from aggregation (the limitation imposed by commercial E6) to symmetry breaking and induction. Even on plates where aggregation succeeded, the proportion of aggregates that elongated and acquired marker expression varied between independent plates, with outcomes assessable only at 72 h, after the critical induction window had closed (Supplementary Figure S3).

We reasoned that systematic optimisation of the remaining induction parameters — pre-treatment medium, seeding density and CHIR99021 dose for each new cell line — would require a real-time, quantitative readout of mesoderm induction, rather than endpoint immunostaining. Although the RUES2-GLR line carries existing endogenous reporters for TBXT, SOX2 and SOX17 (Martyn et al., 2018), our objective was to reduce the number of reporters required and to help apply the gastruloid protocol to multiple hPSC lines that have not previously been characterised in this context. As TBXT is the earliest marker of mesoderm specification, marking the onset of gastrulation downstream of Wnt activation, we generated a TBXT-2A-EGFP knock-in reporter in BJNhem20 (Patil, Das and Inamdar, 2026), which we used as the primary live-readout system in this study. GFP fluorescence provides a quantitative readout of the induction step that defines successful gastruloid formation.

### Optimising pre-treatment medium and seeding density critically influence TBXT-GFP induction

Although iE6 resolved aggregation-step failures, inter-experimental variability in symmetry breaking and gastruloid formation persisted. We hypothesised that variability in the pre-treatment step — specifically, inconsistent mesoderm induction in Nutristem XF — was the underlying cause. To identify the pre-treatment medium that consistently induces mesoderm in BJNhem20, we compared five different media used in published gastruloid protocols (Moris et al., 2020; Hamazaki et al., 2024; Avni et al., 2023): mTeSR Plus, Nutristem XF, N2B27, Stemline II and iE6. All conditions used 3.5 µM CHIR99021 for 24 h as pretreatment and were tested at the end of this period. Immunofluorescence on the parental BJNhem20 line revealed a clear hierarchy of TBXT expression across media, with iE6 producing the strongest and most uniform signal and mTeSR Plus the weakest (Figure 2A). Quantification in the BJNhem20 TBXT-EGFP reporter line confirmed this hierarchy: the proportion of TBXT-GFP-positive nuclei after pre-treatment was 98.6% in iE6, compared to 73.8% in Nutristem XF, 69.1% in Stemline II, 64.4% in N2B27 and 45.9% in mTeSR Plus (Figure 2B). iE6 was therefore selected as the pre-treatment medium for subsequent experiments, making the protocol simpler and defined, due to the use of the same, protein-free medium throughout the protocol.

**Figure 2.**
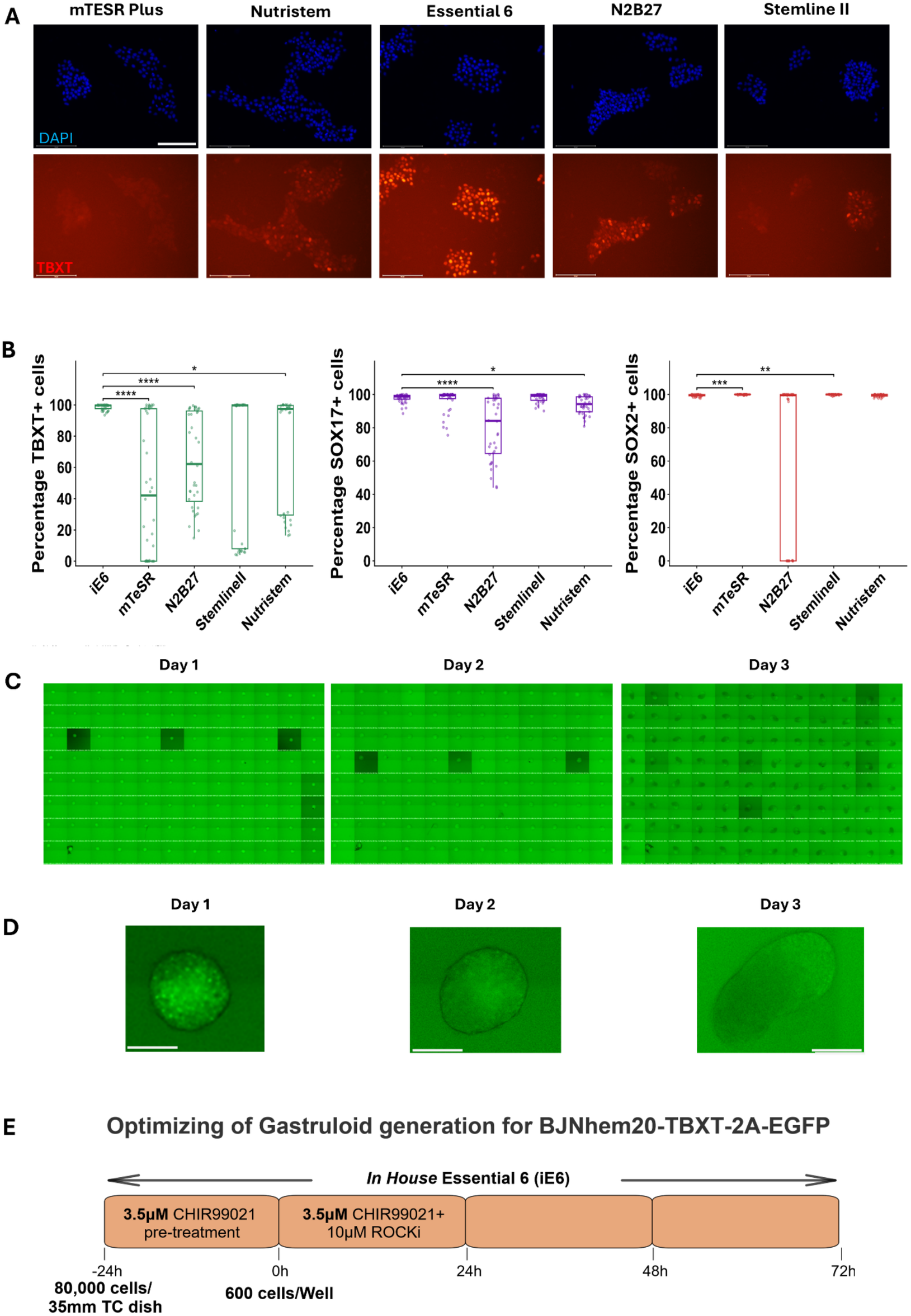
Systematic optimisation of induction parameters reduces inter-experimental variability in gastruloid generation. (A) Immunofluorescence imaging of 2D cultures of BJNhem20 showing expression of TBXT (red) and nuclei (DAPI, blue) following 24-hour CHIR99021 pre-treatment of BJNhem20 cells in five medium formulations: mTeSR Plus, Nutristem XF, in-house Essential 6 (iE6), N2B27, and Stemline II. In-house Essential 6 produced the strongest and most uniform TBXT induction, while the other media gave low or inconsistent expression. Scale bar: 275 µm. (B) Percentage of cells per field of view expressing TBXT, SOX17 or SOX2 post CHIRON induction in different media. Data is represented as box plots showing median (centre line), IQR (whiskers) with individual datapoints overlaid. (N=3; n=34-36). (C) Whole 96-well plate TBXT-EGFP fluorescence images of BJNhem20 TBXT-EGFP gastruloids at days 1, 2 and 3 post-aggregations, seeded at 80,000 cells per 35 mm dish during pre-treatment. (D) Magnified view of representative individual BJNhem20 TBXT-EGFP gastruloids from plate shown in (C). Scale bar, 275 µm. (E) Schematic of the optimized protocol framework with a shorter timeline for gastruloid generation, protocol used in the rest of the study stopping at 72hrs.

Cell density at the time of CHIR99021 exposure determines monolayer confluency and therefore the effective concentration and spatial distribution of Wnt signalling experienced by each cell (Heemskerk et al., 2019). Reproducible gastruloid formation therefore requires that confluency at the start of pre-treatment is tightly controlled, and this in turn requires uniform single-cell suspensions at seeding. We accordingly calibrated the starting conditions by seeding single-cell suspensions of BJNhem20 TBXT-EGFP at defined densities, allowing 24 h for attachment, and then quantified TBXT-GFP expression after the subsequent CHIR99021 pre-treatment to identify the seeding density that produced the most consistent induction.

Cells seeded at densities of 20,000, 40,000, 80,000 and 100,000 per 35 mm tissue culture dish, reached the confluency that resulted in successful gastruloids with polarised TBXT-GFP expression and >90% elongation efficiency in 5, 4 or 1 day (for both 80,000 and 100,000) respectively. Seeding at 80,000 cells per dish for pre-treatment, was adopted for all subsequent experiments as the outcome was consistent with >90% of typical gastruloids generated in at least three independent experiments (Figure 2B) (Supplementary Figure S6; Supplementary Table S4).

### Reproducible single-cell transcriptomic profiles across independently generated gastruloids

To characterize the cellular composition of gastruloids generated using the optimized protocol and to assess transcriptional reproducibility across independent experiments, we performed single-cell RNA sequencing (scRNA-seq) on BJNhem20 TBXT-EGFP gastruloids at 72 hours post-aggregation- the timepoint at which aggregates consistently display axial elongation and germ-layer marker polarisation (Figure 1E). Three individual gastruloids were selected for sequencing, each originating from a separate 96-well plate that had been independently established using cells from distinct 35 mm pre-treatment dishes (Dish1_E5, Dish2_B5 and Dish3_B10). This design enabled simultaneous assessment of transcriptional composition between individual gastruloids and reproducibility across independent biological replicates. A fourth library was generated from a pooled sample containing three additional gastruloids — one drawn from each of the three independent experiments — and processed in parallel as an internal cross-check that cells mix across plates independent of dish of origin; full results from the pooled sample are presented in Supplementary Figure S5. Initial analysis of the dataset was performed using standard log normalization without prior gene filtering, following the canonical Seurat workflow (NormalizeData, FindVariableFeatures, ScaleData). This analysis produced a consistent transcriptional landscape across the three individual gastruloids, identifying 17 unsupervised clusters that resolved into 17 annotated cell populations, including mesodermal, neural and progenitor states broadly aligned with those reported in Moris et al. (2020) (Supplementary Figure S7). To improve the rigor and comparability of the analysis, the dataset was subsequently reprocessed using the analytical framework described by Hamazaki et al. (2024) for scRNA-seq of human gastruloids. This approach applies SCTransform normalization, which corrects for sequencing depth effects and stabilizes variance across genes and incorporates a gene filtering step based on criteria defined by Hamazaki et al. (2024), with low-quality or uninformative genes excluded prior to dimensionality reduction. The annotated lineage populations and their relative proportions were broadly consistent between the two analytical approaches, supporting that the observed cell type composition reflects true biological signal rather than an artifact of the normalization strategy. All results described below and shown in Figure 3 are based on this analysis.

**Figure 3.**
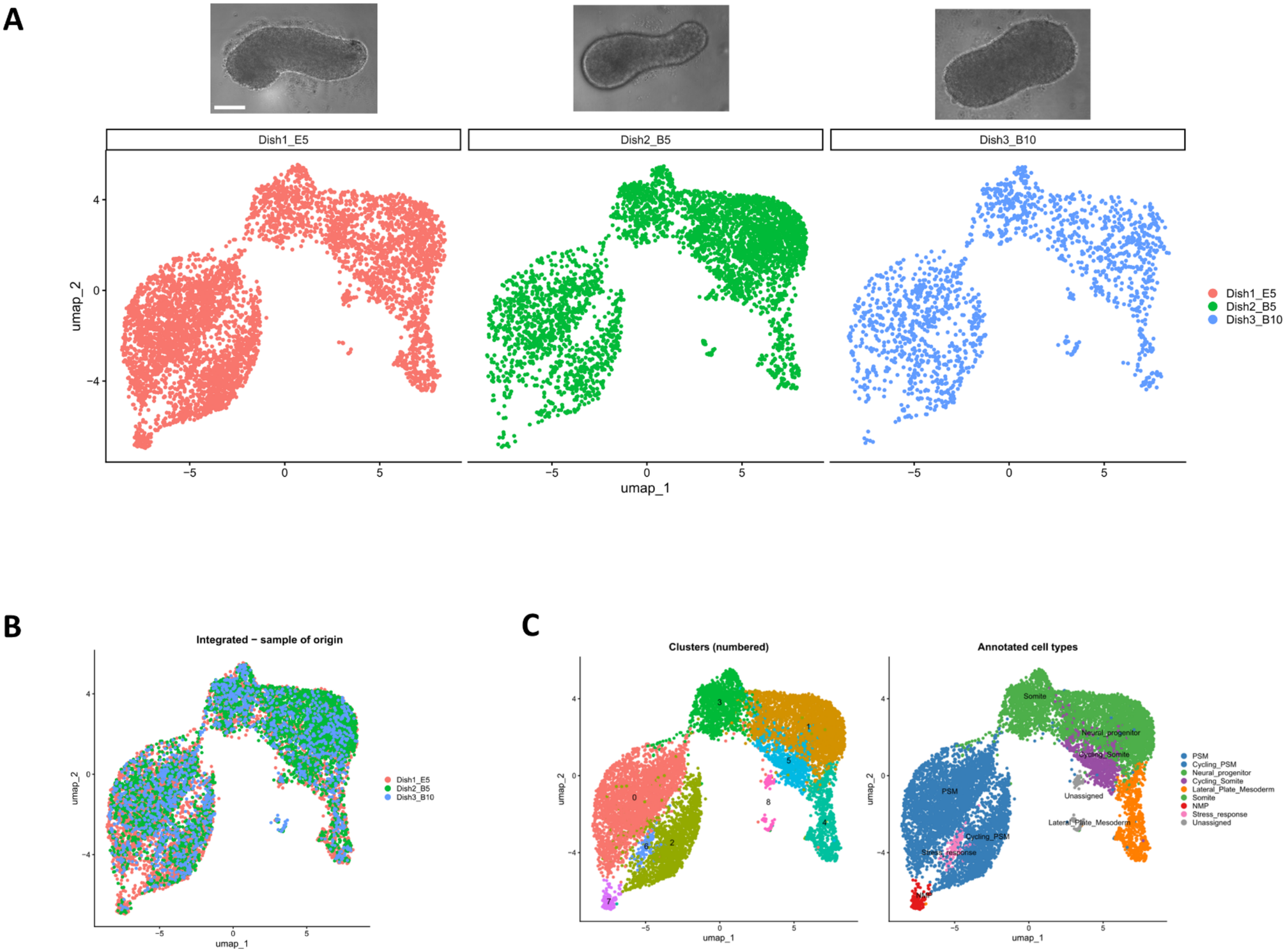
Single-cell transcriptomic profiling reveals reproducible lineage composition across independently generated gastruloids. (A) UMAP embeddings of the Harmony-integrated dataset displayed separately for each of the three independently generated and individually sequenced BJNhem20 TBXT-EGFP gastruloids (Dish1_E5, Dish2_B5, Dish3_B10), with cells coloured by dish of origin. Phase contrast images of the gastruloids used are shown above the respective UMAP. All three samples populate the same UMAP regions, indicating that the same cell types are present across biological replicates. (B) Integrated UMAP showing all cells from the three individually sequenced gastruloids simultaneously, coloured by sample of origin (Dish1_E5, pink; Dish2_B5, green; Dish3_B10, blue). Cells from all three samples are broadly co-distributed across the embedding, confirming that Harmony batch correction successfully aligned the three replicates without collapsing biological variation. (C) Left: integrated UMAP coloured by unsupervised Seurat cluster (clusters 0–8). Right: same UMAP coloured by annotated cell-type identity. Annotated populations include presomitic mesoderm (PSM), somite, lateral plate mesoderm, neuromesodermal progenitors (NMPs), neural progenitors, primitive streak/early mesoderm, a stress-response cluster, and one unassigned population.

Following quality control filtering, SCTransform normalization and Harmony-based batch correction, the three individual gastruloid transcriptomes were integrated into a shared low-dimensional embedding. To assess whether cells from each independently generated gastruloid occupied the same transcriptional space, the integrated Uniform Manifold Approximation and Projection (UMAP) was visualized separately for each gastruloid, with cells colored by dish of origin (Figure 3A). Cells from Dish1_E5, Dish2_B5 and Dish3_B10 distributed across all the major regions of the embedding, with no sample confined to a unique region; differences in absolute cell number across samples reflect differences in cells captured at sequencing rather than sample-specific cell-type composition.

To further visualize the extent of transcriptional overlap, the integrated UMAP was rendered with all cells displayed simultaneously and colored by sample of origin (Figure 3B). Cells from Dish1_E5, Dish2_B5and Dish3_B10 were broadly co-distributed across the embedding, with no transcriptional cluster dominated by cells from a single gastruloid. The three independently generated gastruloids therefore share the same cell-type composition, supporting reproducibility of the optimised protocol at the single-cell level.

Unsupervised clustering of the integrated dataset identified nine transcriptionally distinct clusters (clusters 0–8; Figure 3C, left). Cell type annotation resolved these clusters into eight named populations and one unassigned population (Figure 3C, right). Cluster annotations were assigned by combining marker-gene expression with reference-based classification using SingleR. The annotated populations comprise presomitic mesoderm (PSM), representing the dominant cell population and occupying the largest region of the UMAP; Cycling_PSM, a proliferative subpopulation within the PSM; Somite, reflecting cells progressing along the somitogenesis axis; Cycling_Somite, a cycling subpopulation within the somite compartment; Lateral_Plate_Mesoderm; Neural_progenitor, occupying a distinct region of the embedding consistent with ectodermal and neural specification; and Neuromesodermal Progenitor (NMP), a small population positioned at the boundary between the PSM and neural progenitor compartments, occupying a position consistent with NMP identity at equivalent developmental stages (Hamazaki et al., 2024; Moris et al., 2020).

A Stress_response cluster was identified and retained in the analysis. Stress signatures are commonly observed in scRNA-seq data derived from 3D aggregates and most likely reflect a hypoxic or mechanically stressed core population rather than a biologically distinct lineage—a documented artifact of 3D culture systems reported in prior gastruloid and organoid scRNA-seq datasets (Bhaduri et al., 2020; López-Anguita et al., 2022; Vértesy et al., 2022; Yi et al., 2025). No filtering was applied to this population in the current analysis.

The lineage populations identified here — PSM, somite, lateral plate mesoderm, NMP and neural progenitor — span the major mesodermal and neuroectodermal compartments expected of human gastruloids at this stage. The reproducible distribution of cells from all three gastruloids across shared transcriptional clusters validates the optimised protocol at the single-cell level.

### Validating the protocol in RUES2-GLR and BJNhem19 hPSC

To evaluate whether the optimized gastruloid induction protocol is broadly applicable beyond BJNhem20, we tested its performance in two additional hPSC lines representing distinct genetic backgrounds: the widely used reference line RUES2-GLR (Moris et al., 2020) and the Indian-origin line BJNhem19, a sibling line of BJNhem20. We hypothesised that the core protocol elements — iE6 as the pre-treatment and aggregation medium, 96-well U-bottom aggregation, and pre-treatment to the confluency at which TBXT induction is maximal— would be generally applicable across lines, while the CHIR99021 doses at pre-treatment and pulse, and the cell number at aggregation, would require empirical line-specific titration. Pre-treatment confluency was assessed visually in each line and adjusted based on TBXT levels.

#### RUES-GLR

Empirical titration identified the optimal conditions for RUES2-GLR as 40,000 cells per 35 mm dish for pre-treatment with 3.25 µM CHIR99021, followed by aggregation of 600 cells per well in iE6. CHIR99021 pulse concentrations of 0.5 µM and 1.0 µM, tested along with DMSO vehicle control, produced elongated gastruloids with the characteristic teardrop morphology by 72 h post-aggregation, in three independent experiments (Figure 4A, B). As only the SOX2-mCitrine reporter was compatible with the available imaging system, expression in live aggregates was assessed by mCitrine fluorescence. Polarized mCitrine signal in the expected pattern was observed consistently only with 0.5 µM CHIR99021, hence this was chosen as the effective pulse concentration for RUES2-GLR pulse. Whole-mount immunofluorescence for TBXT, SOX17 and SOX2 at 72 h confirmed a well-defined anteroposterior axis with polarised TBXT enrichment and characteristic spatial restriction of the three markers (Figure 4C). Hence 0.5 μM CHIR99021 was chosen as the effective pulse concentration for RUES2-GLR.

**Figure 4.**
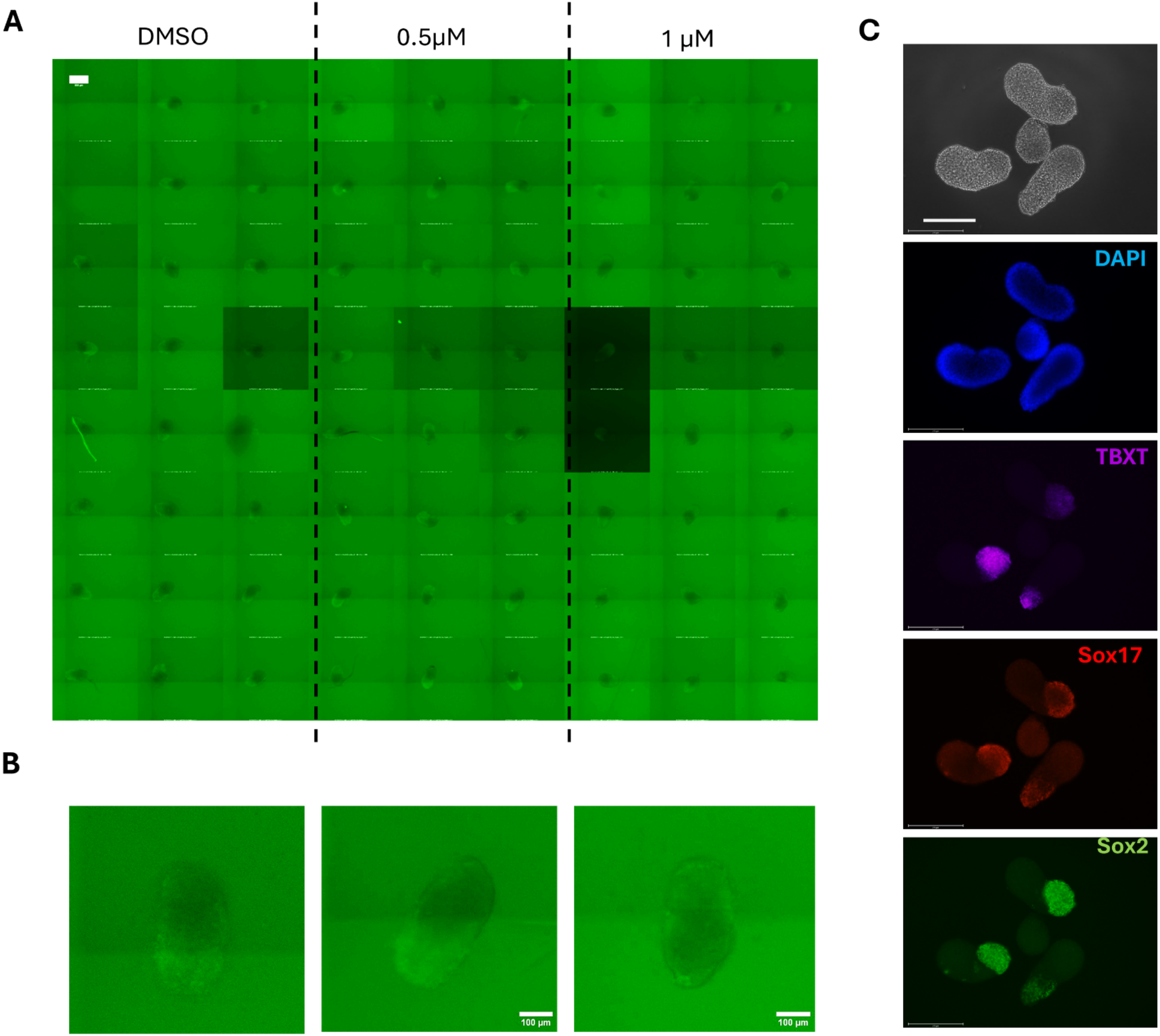
The optimised protocol supports gastruloid formation in RUES2-GLR (panels A–C): (A) Wells from part of a 96-well plate used for generating RUES2-GLR gastruloids, shown at 72 hours post-aggregation. Conditions: 3.25 µM CHIR99021 pre-treatment, with aggregation pulse at DMSO (vehicle), 0.5 µM, and 1.0 µM CHIR99021. 600 cells per well were seeded into U-bottom 96-well plates. (B) Representative individual RUES2-GLR gastruloids at 72 hours post- aggregation from the conditions in (A) (DMSO, 0.5 µM, 1.0 µM CHIR99021 pulse), imaged for endogenous SOX2-mCitrine fluorescence. Polarised Sox2 expression and elongation was observed at all conditions. Scale bar, 100 µm. (C) Whole-mount immunofluorescence of RUES2-GLR gastruloids at 72 hours post-aggregation showing expression of TBXT (magenta), SOX17 (red), and SOX2 (green). Nuclei stained in blue (DAPI). Characteristic polarized expression is seen. Scale bar, 275 µm.

### BJNhem19/BJNhem19-TBXT-2A-EGFP

To enable real-time monitoring of mesoderm induction in this line, we used a BJNhem19 TBXT-EGFP knock-in reporter (hPSCreg JNCSRe001-A-1) as the primary system. Empirical titration identified the optimal conditions as 60,000 cells seeded per 35 mm dish, 2.0 µM CHIR99021 for pre-treatment and 0.5 µM CHIR99021 pulse at aggregation. 200, 400 and 600 cells per well, at aggregation, were tested. 400 cells per well produced the most consistent compact aggregates with reproducible elongation and was adopted for all subsequent experiments (Figure 5A, B). CHIR99021 pulse concentrations of 1.0 and 1.5 µM were also tested in the parental lines and resulted in compact aggregates that failed to elongate (Supplementary Figure S6). Whole-mount immunofluorescence on parental BJNhem19 and BJNhem*19-TBXT-2A-EGFP* gastruloids at 72 h confirmed polarised expression of TBXT, SOX17 and SOX2 (Supplementary S6; Figure 5C), consistent with anteroposterior axis formation and germ layer specification. Two independent experiments yielded comparable results.

**Figure 5.**
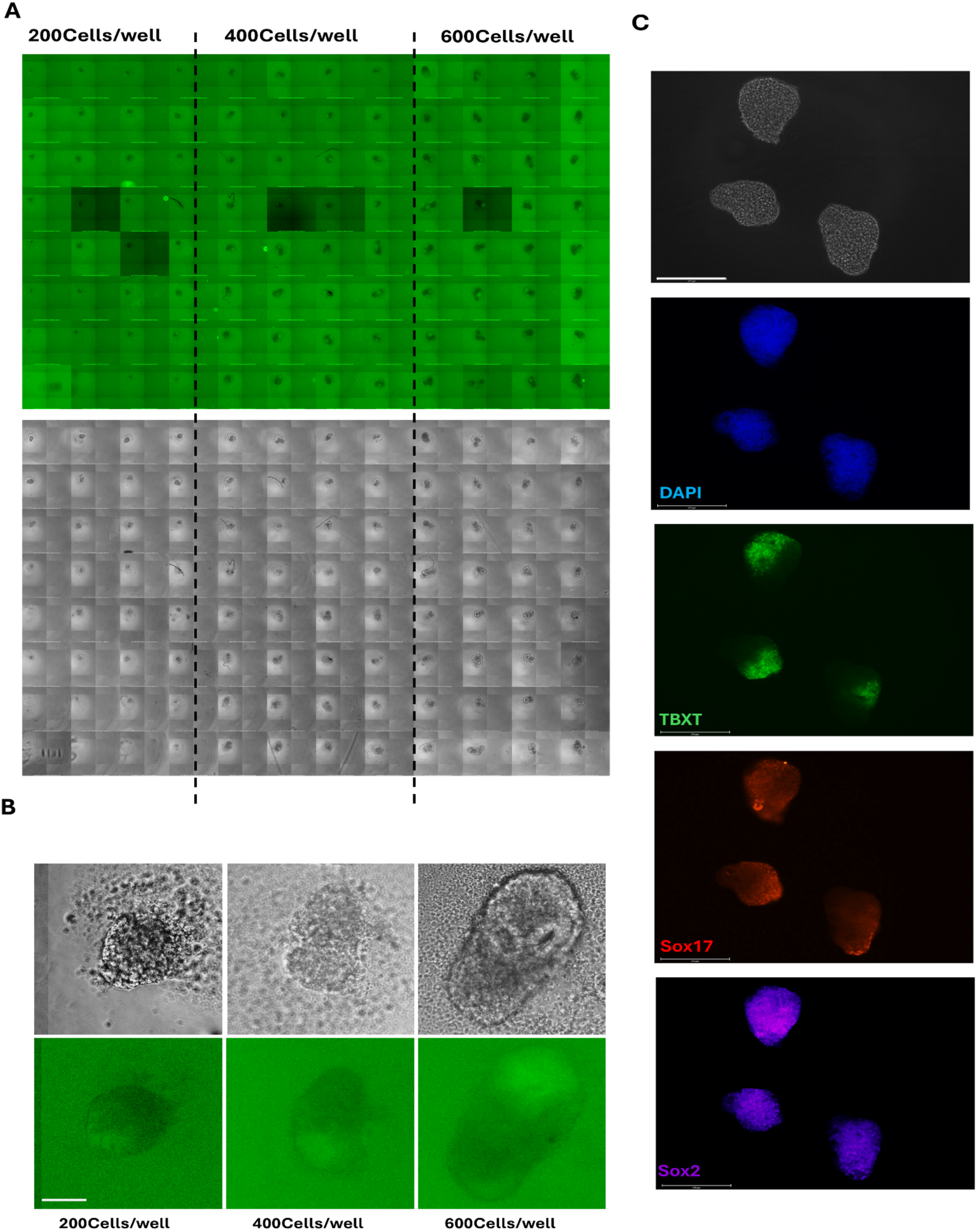
The optimised protocol supports gastruloid formation in BJNhem19 TBXT-EGFP (panels A–C): (A) 96-well plate of BJNhem19 TBXT-EGFP gastruloids at 72 hours post-aggregation. top: TBXT-EGFP fluorescence. bottom: phase-contrast. Conditions: 2.0 µM CHIR99021 pre-treatment with a 0.5 µM aggregation pulse, with founding cell numbers of 200, 400, and 600 cells per well. (B) Magnified view of representative individual BJNhem19 TBXT-EGFP gastruloids from (A), showing TBXT-EGFP fluorescence. Scale bar, 100 µm. (C) Whole-mount immunofluorescence staining of BJNhem19 gastruloids (400 cells per well) at 72 hours, showing expression of TBXT (green), SOX17 (red), SOX2 (magenta). Nuclei stained in blue (DAPI). Scale bar:275 µm.

## Discussion

Human 3D gastruloids (Moris et al., 2020) provide a tractable model of post-implantation development with growing relevance to developmental toxicity screening and disease modelling. However, broad adoption has been constrained by protocol sensitivity to initial conditions, inter-experimental variability and validation in only a small number of hPSC lines — a limitation that is particularly consequential for laboratories with limited resources and access. The work presented here addresses these constraints through a parameter-controlled protocol with three components new to the human gastruloid field: 1] a simple protein-free medium that can be made in-house and used throughout the protocol, namely in-house Essential 6 medium (iE6); 2] Real-time tracking of gastruloid progression based on a single reporter that helps predict gastruloid success and can minimize time to a Go/No Go decision during protocol optimization and 3] validation across three hPSC lines of distinct genetic background — including the previously untested Indian-origin lines BJNhem20 and BJNhem19.

### Defined, in-house medium as a controllable variable

The superior performance of freshly prepared iE6 over four commercial or standard alternatives identifies the basal aggregation medium as a major and previously underappreciated source of inter-experimental variability. A few reports have used freshly prepared medium for gastruloid generation, however, the basal medium is commercially sourced. iE6 is fully defined, serum- and protein-free, prepared from individually sourced components and therefore GMP-compatible. It eliminates reliance on commercial media of unknown composition and makes media quality a controllable parameter, thereby making the protocol globally accessible (Inamdar, 2023).

### A live reporter enables iterative calibration

The BJNhem20 TBXT-2A-EGFP knock-in line addresses a fundamental limitation of fixed-endpoint immunostaining, in which protocol failures become apparent only after the critical induction window has closed and the contributing variables can no longer be resolved. A live transcriptional reporter shortens this feedback cycle, allowing iterative calibration of induction conditions for each experimental configuration or cell line. The reporter is also compatible with automated imaging and high-content screening workflows, lowering the technical barrier for laboratories without prior gastruloid expertise.

### Transferability with line-specific calibration

Validation in BJNhem19 and RUES2-GLR confirms that the core framework — iE6 medium, defined pre-treatment confluency, 96-well aggregation, and a 72-h endpoint — is broadly transferable. CHIR99021 dose at both the pre-treatment and pulse steps varied substantially between lines, however, indicating that Wnt pathway sensitivity must be treated as an empirical, line-specific property that cannot be predicted from genetic background, derivation history or relatedness. Further, as optimal conditions differ for the sibling lines BJNhem19 (male) and BJNhem20 (female) that were derived and maintained under identical conditions, this also raises the possibility of sex-specific differences in protocol response, which can now be investigated. The TBXT-EGFP reporter framework, applicable to any line carrying an equivalent knock-in, supports such line-by-line calibration.

### Limitations

Analysis of the transcriptomes of individual BJNhem20 gastruloids at single-cell resolution, showed high reproducibility. Comprehensive scRNA-seq profiling at multiple timepoints of gastruloid formation and in additional lines, will be required to establish the model’s full biological and translational relevance.

### Applicability

Baseline reproducibility must be established before gastruloid models can be applied in genetic, pharmacological or environmental perturbation-based experimental designs. Gastruloids have great potential for translational applications including developmental toxicity screening (Mantziou et al., 2021), particularly given regulatory shifts in several jurisdictions toward reducing animal-based teratogenicity assays. The over-90% elongation efficiency demonstrated here permits scale-up within single 96-well plates, increasing throughput and statistical power. Combined with the cost reductions of in-house medium and the live reporter for early assessment of experimental outcome, this framework lowers the technical and economic barriers to using gastruloids as a genuinely human-relevant developmental model. Extending validation to hPSC lines from additional under-represented populations is the natural next step.

## Author Contributions

**Arghakusum Das**: Conceptualization, Methodology, Formal analysis, Investigation, Data curation, Visualization, Writing – original draft, Writing – review & editing. **Shekhar Patil:** Methodology, Resources, Data curation. **Karthik Ravi**: Methodology, Data curation **Maneesha S. Inamdar:** Conceptualization, Methodology, Resources, Supervision, Funding acquisition, Writing – Original Draft, review & editing.

## Acknowledgements

We thank Kavin Manikandan for help with IF studies in different culture media, Prof. Alfonso Martinez Arias for discussions and for helpful suggestions and feedback on the manuscript, and members of Inamdar laboratory for valuable discussion and suggestions.

## References

Ai Z, Niu B, Yin Y, Xiang L, Shi G, Duan K, et al. Dissecting peri-implantation development using cultured human embryos and embryo-like assembloids. Cell Res. 2023;33(9):661–678. doi:10.1038/s41422-023-00846-8

Avni, Lara, Naama Farag, Binita Ghoshand Iftach Nachman. “Gastruloid optimization.” Emerging Topics in Life Sciences 7, no. 4 (2023): 409–415. doi:10.1042/ETLS20230096

Aran D, Looney AP, Liu L, Wu E, Fong V, Hsu A, et al. Reference-based analysis of lung single-cell sequencing reveals a transitional profibrotic macrophage. Nat Immunol. 2019;20(2):163–172. doi:10.1038/s41590-018-0276-y

Arias AM, Rivron N, Tajbakhsh S, et al. Human stem cell-based embryo models: innovation, ethicsand policy. Hum Reprod. Published online March 24, 2026. doi:10.1093/humrep/deag035

Beccari L, Moris N, Girgin M, Turner DA, Baillie-Johnson P, Cossy AC, et al. Multi-axial self-organization properties of mouse embryonic stem cells into gastruloids. Nature. 2018;562(7726):272–276. doi:10.1038/s41586-018-0578-0

Bhaduri Aandrews MG, Mancia Leon W, Jung D, Shin D, Allen D, et al. Cell stress in cortical organoids impairs molecular subtype specification. Nature. 2020;578(7793):142–148. doi:10.1038/s41586-020-1962-0

Chen G, Gulbranson DR, Hou Z, Bolin JM, Ruotti V, Probasco MD, et al. Chemically defined conditions for human iPSC derivation and culture. Nat Methods. 2011;8(5):424–429. doi:10.1038/nmeth.1593

Chepda T, Cadau M, Girin P, Frey J, Chamson A. Monitoring of ascorbate at a constant rate in cell culture: effect on cell growth. In Vitro Cell Dev Biol Anim. 2001;37(1):26–30. doi:10.1290/1071-2690(2001)037<0026:MOAAAC>2.0.CO;2

European Parliament and Council of the European Union. Directive 2010/63/EU of the European Parliament and of the Council of 22 September 2010 on the protection of animals used for scientific purposes. Off J Eur Union. 2010;L 276:33-79. https://eur-lex.europa.eu/legal-content/EN/TXT/?uri=CELEX:32010L0063

Hafemeister C, Satija R. Normalization and variance stabilization of single-cell RNA-seq data using regularized negative binomial regression. Genome Biol. 2019;20(1):296. doi:10.1186/s13059-019-1874-1

Hamazaki N, Yang W, Kubo CA, Qiu C, Martin BK, Garge RK, et al. Retinoic acid induces human gastruloids with posterior embryo-like structures. Nat Cell Biol. 2024;26(10):1790–1803. doi:10.1038/s41556-024-01487-8

Heemskerk I, Burt K, Miller M, Chhabra S, Guerra MC, Liu L, Warmflash A. Rapid changes in morphogen concentration control self-organized patterning in human embryonic stem cells. eLife. 2019;8:e40526. doi:10.7554/eLife.40526

Inamdar, Maneesha S. “Yes to global standards for research—as long as they are truly global.” Nature 623, no. 7987 (2023): 460–460. doi:10.1038/d41586-023-03445-2

Inamdar MS, Venu P, Srinivas MS, Rao K, VijayRaghavan K. Derivation and characterization of two sibling human embryonic stem cell lines from discarded grade III embryos. Stem Cells Dev. 2009;18(3):423–433. doi:10.1089/scd.2008.0131

Itskovitz-Eldor J, Schuldiner M, Karsenti D, Eden A, Yanuka O, Amit M, Soreq H, Benvenisty N. Differentiation of human embryonic stem cells into embryoid bodies comprising the three embryonic germ layers. Mol Med. 2000;6(2):88–95. doi:10.1007/BF03401776

Korsunsky I, Millard N, Fan J, Slowikowski K, Zhang F, Wei K, et al. Fast, sensitive and accurate integration of single-cell data with Harmony. Nat Methods. 2019;16(12):1289–1296. doi:10.1038/s41592-019-0619-0

Liu L, Oura S, Markham Z, Hamilton JN, Skory RM, Li L, et al. Modeling post-implantation stages of human development into early organogenesis with stem-cell-derived peri-gastruloids. Cell. 2023;186(18):3776–3792.e16. doi:10.1016/j.cell.2023.07.018

López-Anguita N, Gassaloglu SI, Stötzel M, Bolondi A, Conkar D, Typou M, et al. Hypoxia induces an early primitive streak signature, enhancing spontaneous elongation and lineage representation in gastruloids. Development. 2022;149(20):dev200679. doi:10.1242/dev.200679

Makwana K, Tilley L, Chakravarty P, et al. Modelling co-development between the somites and neural tube in human trunk-like structures. Nat Cell Biol. 2025;27(12):2049–2062. doi:10.1038/s41556-025-01813-8

Mantziou, Veatriki, Peter Baillie-Benson, Marloes Jaklin, Stefan Kustermann, Alfonso Martinez Ariasand Naomi Moris. “In vitro teratogenicity testing using a 3D, embryo-like gastruloid system.” Reproductive Toxicology 105 (2021): 72–90. doi:10.1016/j.reprotox.2021.08.003

McInnes L, Healy J, Melville J. UMAP: Uniform Manifold Approximation and Projection for dimension reduction. J Open Source Softw. 2018;3(29):861. doi:10.21105/joss.00861

Michels AJ, Frei B. Myths, artifactsand fatal flaws: identifying limitations and opportunities in vitamin C research. Nutrients. 2013;5(12):5161–5192. doi:10.3390/nu5125161

Moris N, Anlas K, Schröder J, Ghimire S, Balayo T, Martinez Arias A, et al. Generating human gastruloids from human embryonic stem cells. Protocol Exchange. 2020. doi:10.21203/rs.3.pex-812/v1

Moris N, Anlas K, van den Brink SC, Alemany A, Schröder J, Ghimire S, et al. An in vitro model of early anteroposterior organization during human development. Nature. 2020;582(7812):410–415. doi:10.1038/s41586-020-2383-9

Patil S, Das A, Inamdar MS. Generation of a Brachyury reporter cell line (BJNhem20 Brachyury (TBXT)-2A-EGFP) in human embryonic stem cells using CRISPR-Cas9 gene targeting. Stem Cell Res. 2026;91:103907. doi:10.1016/j.scr.2026.103907

Rivron NC, Frias-Aldeguer J, Vrij EJ, Boisset JC, Korving J, Vivie J, et al. Blastocyst-like structures generated solely from stem cells. Nature. 2018;557(7703):106–111. doi:10.1038/s41586-018-0051-0

Schindelin J, Arganda-Carreras I, Frise E, Kaynig V, Longair M, Pietzsch T, et al. Fiji: an open-source platform for biological-image analysis. Nat Methods. 2012;9(7):676–682. doi:10.1038/nmeth.2019

Stuart T, Butler A, Hoffman P, Hafemeister C, Papalexi E, Mauck WM, et al. Comprehensive integration of single-cell data. Cell. 2019;177(7):1888–1902.e21. doi:10.1016/j.cell.2019.05.031

Umans BD, Gilad Y. Oxygen-induced stress reveals context-specific gene regulatory effects in human brain organoids. Genome Res. 2025;35(8):1689–1700. doi:10.1101/gr.280219.124

United States Congress. FDA Modernization Act 2.0, Pub. L. No. 117-328, §3209, 136 Stat. 4459 (2022). https://www.congress.gov/bill/117th-congress/senate-bill/5002

van den Brink SC, Baillie-Johnson P, Balayo T, Hadjantonakis AK, Nowotschin S, Turner DA, Martinez Arias A. Symmetry breaking, germ layer specification and axial organisation in aggregates of mouse embryonic stem cells. Development. 2014;141(22):4231–4242. doi:10.1242/dev.113001

van den Brink SC, Sage F, Vertesy A, Spanjaard B, Peterson-Maduro J, Baron CS, et al. Single-cell sequencing reveals dissociation-induced gene expression in tissue subpopulations. Nat Methods. 2017;14(10):935–936. doi:10.1038/nmeth.4437

Venu P, Chakraborty S, Inamdar MS. Analysis of long-term culture properties and pluripotent character of two sibling human embryonic stem cell lines derived from discarded embryos. In Vitro Cell Dev Biol Anim. 2010;46(3-4):200–205. doi:10.1007/s11626-010-9277-3

Vértesy Á, Eichmüller OL, Naas J, Novatchkova M, Esk C, Balmaña M, oet al. Gruffi: an algorithm for computational removal of stressed cells from brain organoid transcriptomic datasets. EMBO J. 2022;41(17):e111118. doi:10.15252/embj.2022111118

Warmflash A, Sorre B, Etoc F, Siggia ED, Brivanlou AH. A method to recapitulate early embryonic spatial patterning in human embryonic stem cells. Nat Methods. 2014;11(8):847–854. doi:10.1038/nmeth.3016

Yi S, Huang M, Xian C, Kong X, Yin S, Peng J, et al. Single-cell transcriptomics of vascularized human brain organoids decipher lineage-specific stress adaptation in fetal hypoxia-reoxygenation injury. Theranostics. 2025;15(14):7001–7024. doi:10.7150/thno.117001

Yu L, Wei Y, Duan J, Schmitz DA, Sakurai M, Wang L, et al. Blastocyst-like structures generated from human pluripotent stem cells. Nature. 2021;591(7851):620–626. doi:10.1038/s41586-021-03356-y

