## Supplementary Information for "A robust, reproducible, accessible and scalable protocol for generating three-dimensional human gastruloids"

**Supplementary Tables**

**Supplementary Table S1.** Gastruloid elongation efficiency as a function of founding cell number per well (BJNhem20). Elongated gastruloids were defined by visual assessment of aspect ratio at 72 hours post-aggregation.

| Cells per well | Independent experiments (n) | Wells tested | Elongated gastruloids | Efficiency (%) |
|---|---|---|---|---|
| **400** | 4 | 144 | 132 | 91.7 |
| **500** | 1 | 48 | 4 | 8.3 |
| **600** | 6 | 192 | 180 | 93.8 |
| **800** | 2 | 48 | 36 | 75.0 |
| **1000** | 2 | 12 | 0 | 0.0 |

**Supplementary Table S2.** Plate-level outcomes from 20 independent runs of the optimised gastruloid protocol in BJNhem20 using iE6 as the aggregation medium.

| Plate | Wells with elongated gastruloids / total wells | Elongation Outcome |
|---|---|---|
| **1** | 96/96 | Success |
| **2** | 0/96 | Failed |

| | | |
|---|---|---|
| **3** | 0/96 | Failed |
| **4** | 41/96 | Success |
| **5** | 84/96 | Success |
| **6** | 48/48 | Success |
| **7** | 0/48 | Failed |
| **8** | 19/48 | Success |
| **9** | 0/48 | Failed |
| **10** | 41/47 | Success |
| **11** | 0/24 | Failed |
| **12** | 41/96 | Success |
| **13** | 0/96 | Failed |
| **14** | 0/24 | Failed |
| **15** | 0/96 | Failed |
| **16** | 0/96 | Failed |
| **17** | 0/96 | Failed |
| **18** | 0/96 | Failed |
| **19** | 96/96 | Success |
| **20** | 72/96 | Success |

**Supplementary Table S3.** Quantification of germ layer marker expression and spatial patterning by whole-mount immunofluorescence in BJNhem20 gastruloids at 72 hours post-aggregation.

| Germ layer marker | Gastruloids positive (n) | Showing correct spatial patterning (n) |
|---|---|---|
| **TBXT/Brachyury (mesoderm)** | 44 | 44 |
| **SOX2 (ectoderm)** | 28 | 28 |
| **SOX17 (endoderm)** | 16 | 16 |

**Supplementary Table S4.** Effect of pre-treatment seeding density on protocol duration and gastruloid formation in BJNhem20 TBXT-EGFP cells. Values are shown as successful/total independent experiments.

| Cells seeded per 35 mm dish | Time to complete protocol (days) | Gastruloid elongation (experiments) | TBXT-EGFP expression (experiments) |
|---|---|---|---|
| **20,000** | 9 | 1/1 | 0/1 |
| **40,000** | 8 | 2/3 | 2/3 |
| **50,000** | 7 | 1/2 | 1/2 |
| **60,000** | 7 | 1/2 | 1/2 |

| | | | |
|---|---|---|---|
| **80,000** | 5–6 | 4/5 | 4/5 |
| **100,000** | 5–6 | 1/2 | 1/2 |

**Supplementary Figures and Legends**

**Supplementary Figure S1**

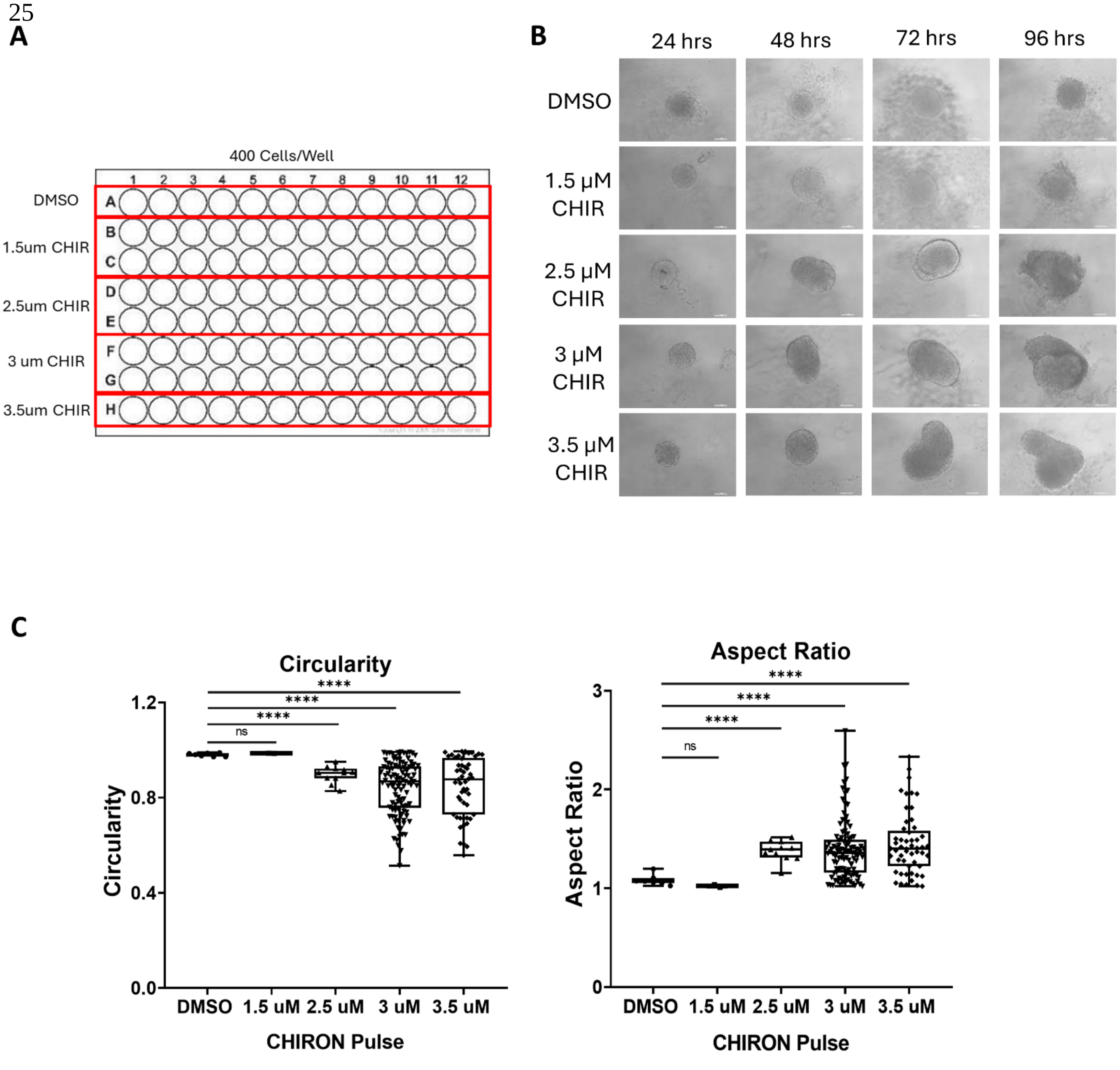

**Supplementary Figure S2**

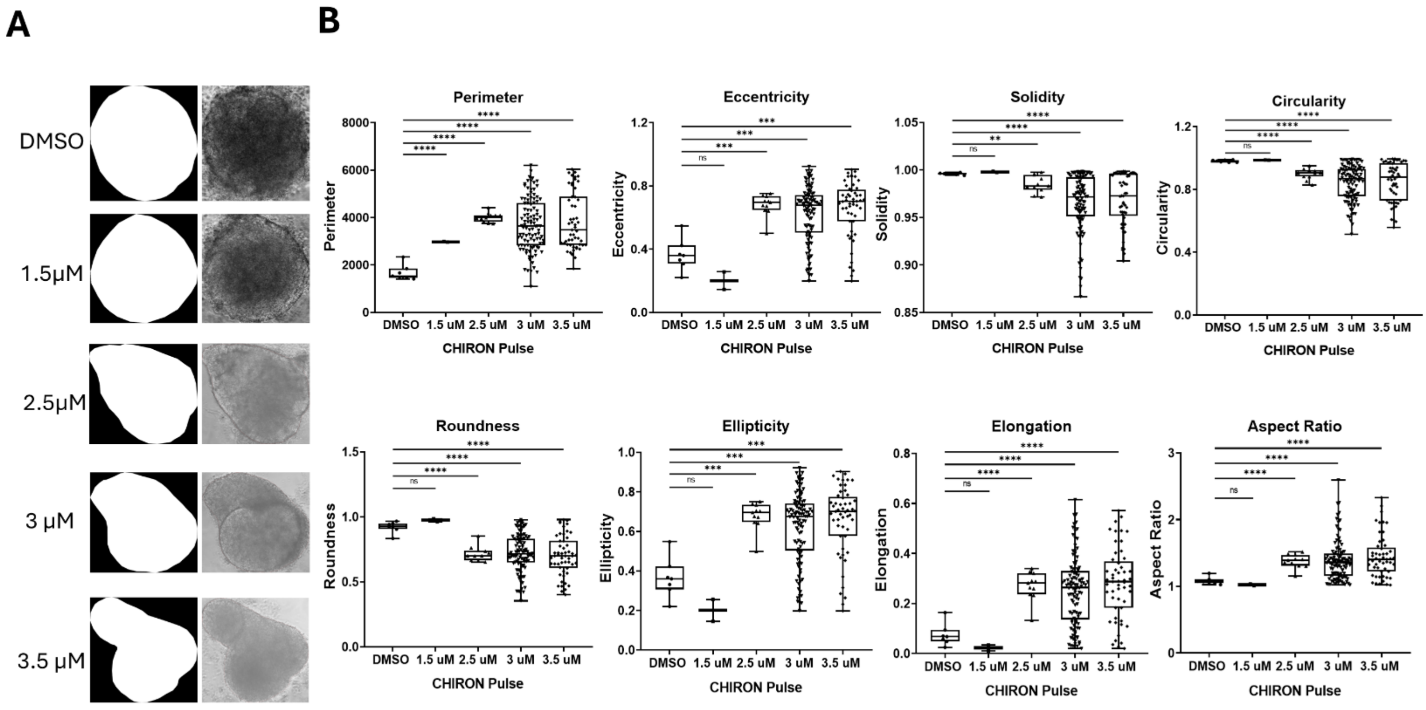

**Supplementary Figure S3**

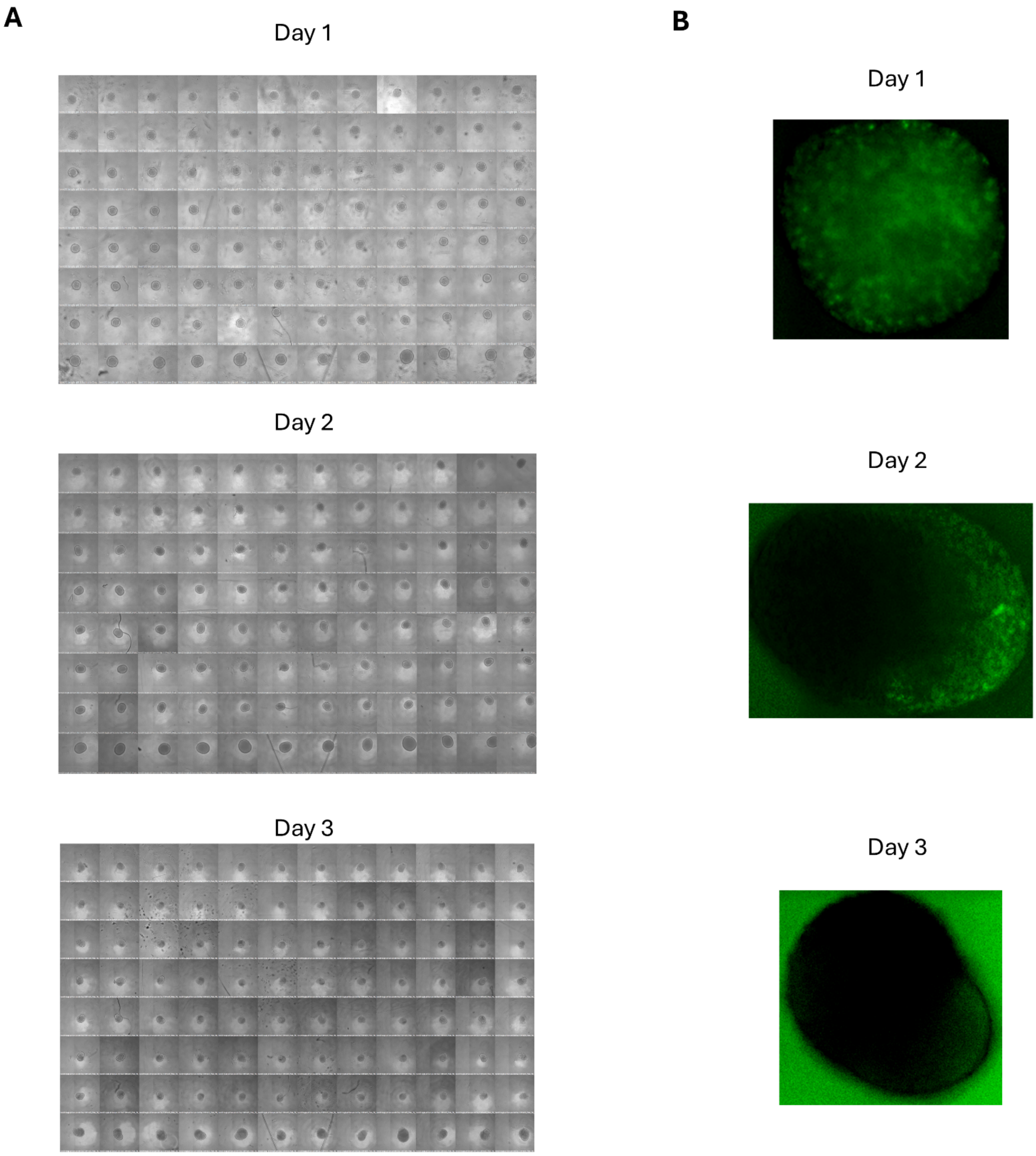

### Supplementary Figure S4

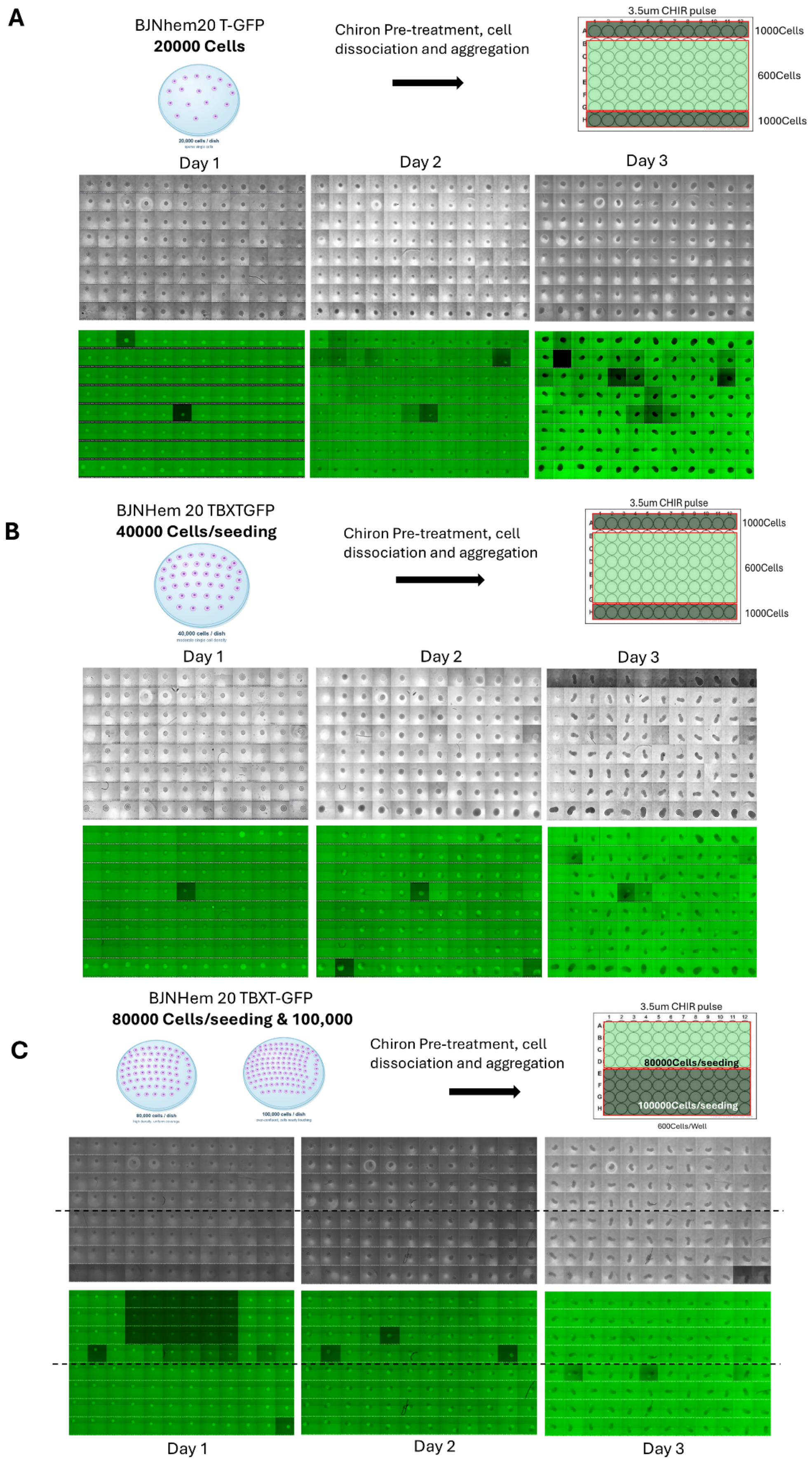

### Supplementary Figure S5

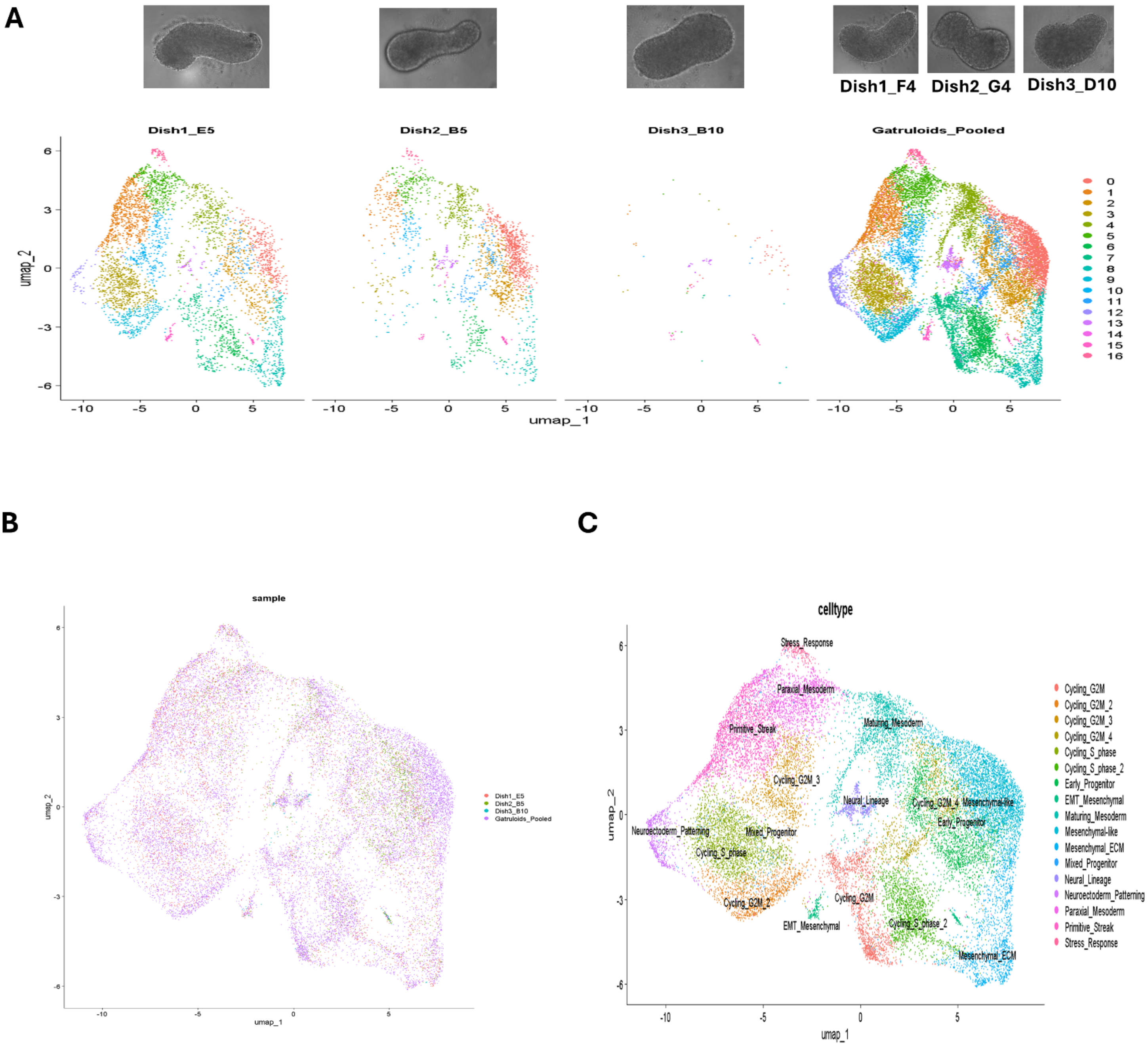

**Supplementary Figure S6**

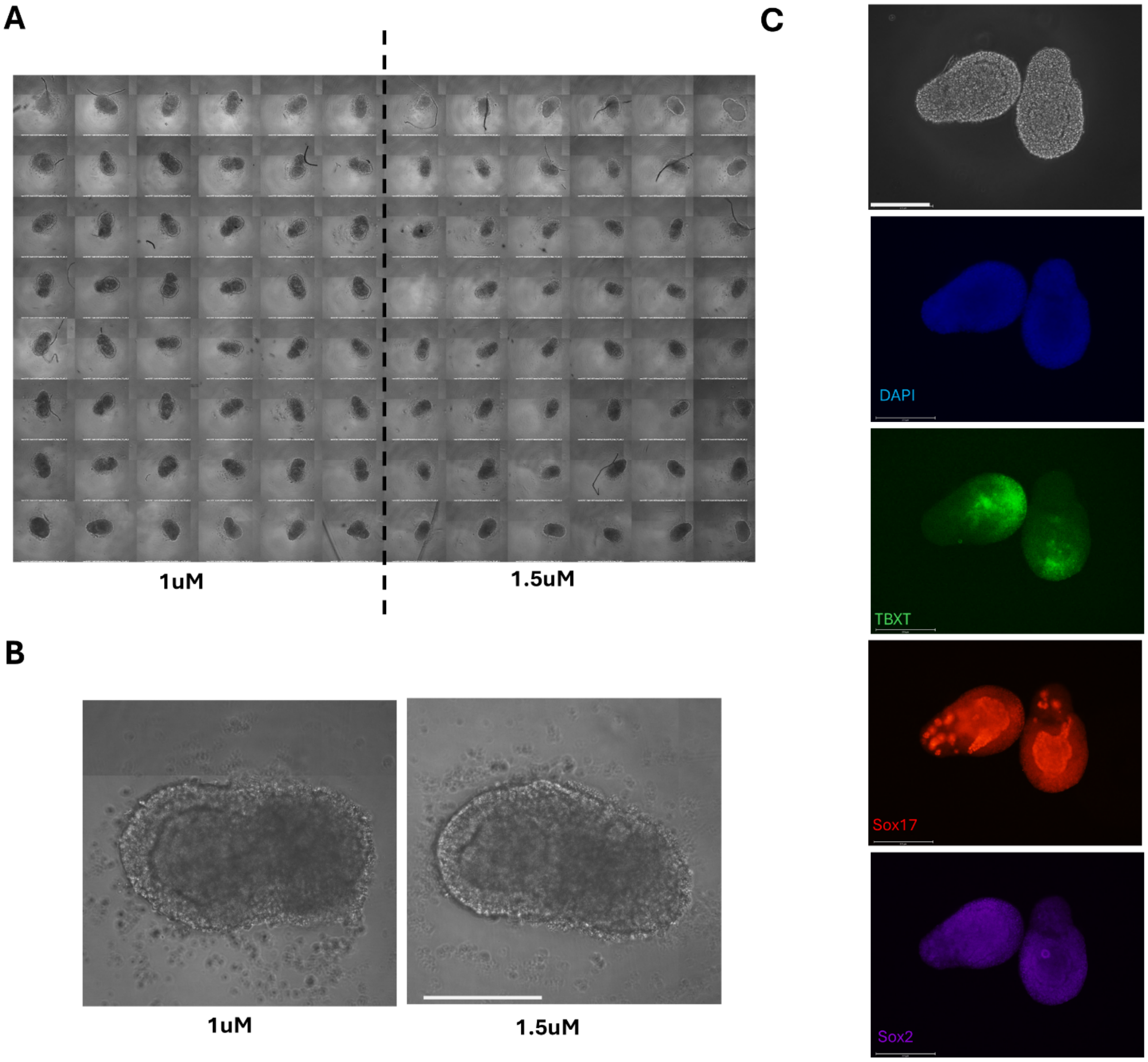

**Supplementary Figure S1. Optimisation of the gastruloid generation protocol in BJNhem20.**

(A) Plate layout for the CHIR99021 pulse titration. 96-well U-bottom plates seeded with 400 BJNhem20 cells per well were exposed to one of five conditions during the 0–24 h pulse window: vehicle control (0.02% DMSO; row A), 1.5 µM CHIR99021 (rows B–C), 2.5 µM (rows D–E), 3.0 µM (rows F–G) or 3.5 µM (row H).

(B) Phase-contrast imaging of representative BJNhem20 aggregates at 24, 48, 72 and 96 h post-aggregation under each CHIR99021 pulse condition shown in (A). All aggregates were spherical at 24 h. By 48 h, symmetry breaking was observed at concentrations of 2.5 µM and above. By 72 h, 3.0 and 3.5 µM produced well-elongated gastruloids with two morphologically distinct poles, whereas 2.5 µM produced less elongated aggregates with protrusions and uneven edges, and 1.5 µM remained indistinguishable from DMSO controls. No further elongation was observed between 72 and 96 h. Scale bar = 100 µm.

(C) Quantitative morphometric analysis of BJNhem20 aggregates at 72 h post-aggregation across the CHIR99021 pulse conditions shown in (A). Left: circularity (1.0 = perfect circle); right: aspect ratio (1.0 = equal axes).

**Supplementary Figure S2. MATLAB-based image-analysis pipeline for extracting morphometric parameters from phase-contrast images.**

Representative workflow showing phase-contrast images of a BJNhem20 gastruloids, the segmented outline produced by polygon-based manual annotation (A), and the resulting morphometric measurements (B), including area, perimeter, circularity, and aspect ratio. The pipeline was adapted from Moris et al. (2020) and implemented in MATLAB R2023a.

**Supplementary Figure S3. Impact of CHIR99021 pre-treatment concentration on gastruloid polarisation**

Phase-contrast images of 96 Well plate (A) and magnified insets of TBXT-EGFP fluorescence images (B) of BJNhem20 TBXT-EGFP gastruloids at days 1, 2, and 3 post-aggregations. Cells were pre-treated with 5.0 μM CHIR99021, resulting in compact, non-elongating aggregates where TBXT-EGFP signal was diffused or absent.

**Supplementary Figure S4. Optimisation of pre-treatment seeding density for TBXT-EGFP expression in BJNhem20 cells.**

BJNhem20 TBXT-EGFP cells were seeded at (A) 20,000, (B) 40,000 and (C) 80,000 & 100,000 cells per 35 mm dish for CHIR99021 pre-treatment, then aggregated and imaged at days 1, 2, and 3 post-aggregations. Phase-contrast (upper rows) and TBXT-EGFP fluorescence (lower rows) are shown; plate layouts are included at right.

**Supplementary Figure S5. Initial scRNA-seq analysis of BJNhem20 TBXT-EGFP gastruloids using standard log normalization identifies broad transcriptional clusters consistent with SCTransform-based analysis.**

(A) Phase-contrast images of individually sequenced gastruloids (Dish1_E5, Dish2_B5, Dish3_B10) and the pooled sample (Gastruloids_Pooled) at 72 hours post-aggregation (top), with corresponding UMAP embeddings from initial log normalization analysis without prior gene filtering, colored by Seurat cluster identity (bottom). Seventeen transcriptionally distinct clusters are identified across the four libraries. (B) Integrated UMAP with all cells colored by sample of origin, showing co-distribution of cells from all three individual gastruloids and the pooled sample across shared clusters. (C) Same UMAP colored by annotated cell type identity from the initial log normalization analysis. The broad lineage composition—including presomitic mesoderm, neuromesodermal progenitor, and neural progenitor states—is concordant with the refined annotation obtained by SCTransform-based analysis shown in Figure 3.

**Supplementary Figure S6. Optimisation of CHIR99021 pulse concentration for BJNhem19 gastruloid formation.**

(A) 96-well plate phase-contrast view of BJNhem19 gastruloids at 72 hours post-aggregation, generated with CHIR99021 pulse concentrations of 1.0 µM and 1.5 µM as indicated. (B) Representative magnified phase-contrast images of individual gastruloids from the 1.0 µM and 1.5 µM pulse conditions, illustrating aggregate morphology and extent of axial elongation at 72 hours. (C)Whole-mount immunofluorescence of BJNhem19 gastruloids generated with the 1.0 µM pulse, fixed at 72 hours and stained for TBXT (green), SOX17 (red), SOX2 (magenta), and DAPI (blue). Scale bar: 275 µm.
